# Resolving the molecular niche of pulmonary fibrosis using cryopreserved human precision cut lung slices

**DOI:** 10.1101/2025.04.22.649643

**Authors:** Indu Sivankutty, Maniteja Arava, Jae Hun Kim, Zheming An, Erica Matsui, Debesh Sahu, Chunhui Cai, Holger Behrsing, Kaia Gerlovin, Matthew Paul Lech, Béla Suki, Ramaswamy Krishnan, Sangita Choudhury

## Abstract

**Background and methods:** Pulmonary fibrosis (PF) manifests locally and heterogeneously in the diseased lung. To uncover mechanistic underpinnings, we have utilized cryopreserved fibrotic human precision-cut lung slices (PCLS), and using single-cell spatial transcriptomics, mapped and interpreted the transcriptomic regulation of effector cell types/states.

**Results:** Cryopreserved-thawed fibrotic human PCLS retained canonical molecular and cellular PF hallmarks including fibroblast and myofibroblast accumulation, epithelial depletion, and vascular remodeling. Notably, these cellular changes were spatially co-incident with 1) collagen-rich regions enriched for extracellular matrix (ECM)-related genes (e.g., COL1A1, SPARC), 2) expanded fibroblast-endothelial crosstalk with enhanced chemokine signaling (CXCL12– CXCR4), 3) integrin-driven mesenchymal signaling targets (ANGPTL–integrin), 4) stress-adapted and immune-interacting Alveolar Type 2 cells with impaired surfactant gene expression, and 5) endothelial reprogramming toward a contractile and profibrotic phenotype. At the transcriptomic level, the spatially-defined fibrotic niche was characterized by increased gene expression related to immune responses (IGKC and IGHG1), fibrosis (collagen, fibronectin, and DCN), mechanotransduction (MYH11, TPM2, ACTG2, ACTA2), immune cell recruitment (CCL2, IL6, ICAM1), ECM remodeling (e.g. thrombospondin 1, THBS1), and decreased gene expression related to surfactant production (SFTPB, SFTPC). At the functional level, the fibrotic niches were defined by the activation of three fibroblast mediating signaling pathways: (1) the matricellular glycoprotein, SPARC, (2) Relaxin signaling, and (3) multiple immunoglobulin transcripts, including IGHA1, IGHG1, IGHG3, and IGKC.

**Conclusions:** We verify model fidelity, identify both known and unanticipated pro-fibrotic genes, and advocate for the expanded use of spatial transcriptomic measurements using cryopreserved human PCLS for disease modeling and drug discovery.

## Background

Fibrotic drug discovery requires preclinical testing using cell-, tissue- or animal-based models. While cell- and tissue-based models including 2D culture, 3D organoids, reconstructed tissues and lung micro-physiological systems ^1–4^ are fast, and permit direct and unambiguous assessment of drug activity, they lack physical, biological and biochemical interactions provided by molecular, intercellular and extracellular interactions of the fibrotic lung. The outcome is missed or false discovery. The animal model more accurately reproduces lung complexity but inference of drug outcomes is slow, time-consuming, and complex. Moreover, due to immunological differences, animal studies cannot fully recapitulate human fibrotic outcomes ^5^. Thus, academic and pharmaceutical leaders have emphasized the need to develop new pre-clinical models that closely resemble the human lung ^6–9^. To this end, thin-cut slices of fibrotic human lungs, called human precision-cut lung slices (PCLS), have become invaluable^10^.

Fibrotic human PCLS are viable, functional, and architecturally intact. Native cells, ECM, and the biochemical niche are largely preserved and interact in realistic ways. The fibrotic PCLS recapitulates excessive collagen deposition, fibroblast-to-myofibroblast differentiation, alveolar epithelial reprogramming ^11^, spatio-temporal heterogeneity of incidence and composition of localized fibrotic lesions ^12^, and pro-fibrotic secretion ^13^. Given these advantages, the fibrotic human PCLS have emerged as a preferred platform for drug testing, genetic manipulation, toxicity assessment, RNA isolation, functional measurements, and histology ^11,13–19^. Moreover, recent enhancements in PCLS procurement and preparation have enhanced throughput to upwards of thousands of PCLS per human lung. While each slice remains viable and functional for up to a month, the implementation of cryopreservation has enhanced PCLS viability even further, to years ^20^, with minimal impact to mitochondrial integrity, glutathione activity, cell viability, tissue integrity, airway contractile and relaxant responses, toxicological outcomes, cytokine responsiveness, phagocytosis and lymphocyte proliferation ^14,20–23^. Thus, between oversimplified measurements in the cultured cell and inferential measurements using animal models, the human fibrotic PCLS bridges an important translational gap by allowing demographic-based donor-donor comparisons and repeat donor testing.

Here, we ask a fundamental question – to what extent does the cryopreserved human fibrotic PCLS recapitulate cellular diversity and intercellular communication inherent in the never-frozen PF lung? We leveraged recent advancements in single-cell RNA sequencing (scRNA-seq) and spatial transcriptomics and implement these methods to 1) evaluate the transcriptional profiles and spatial dynamics of preserved cells within the cryopreserved fibrotic human PCLS, 2) compare differences between human fibrotic and non-fibrotic cryopreserved PCLS, and 3) evaluate similarities to published spatial transcriptomic measurements in never-frozen PF biopsies.

## Material and methods

### Human PCLS procurement

Cryopreserved PCLS were purchased from non-transplantable de-identified human lungs from the Institute for In Vitro Sciences (Gaithersburg, MD)^14,20^. These lungs were obtained from consenting donors (or next of kin) through accredited procurement agencies, including the International Institute for the Advancement of Medicine (IIAM; Edison, NJ, USA), Novabiosis, Inc. (Durham, NC, USA), or National Disease Research Interchange (Philadelphia, PA, USA). The donor demographics are provided in Table 1. Upon thaw, these slices retain biomass, viability, tissue integrity, and inflammatory markers^14^.

**Table 1.** Donor Demographic Information.

| Donor<br>number | Age | Race | Height<br>(cm) | Weight<br>(kg) | Sex | Lung Health<br>Status |
| --- | --- | --- | --- | --- | --- | --- |
| N1-23 | 66 | Hispanic/Latino | 157 | 61.7 | M | Normal |
| N2-62 | 69 | Caucasian | 175 | 79 | M | Normal |
| N3-25 | 60 | White | 168 | 83.7 | M | Normal |
| D1-57 | 65 | Hispanic | NA | NA | F | IPF/ILD |
| D2-7 | 67 | Caucasian | NA | NA | M | IPF/ILD |
| D3-82 | 61 | Black | NA | NA | F | PF/ILD/PH |

### PCLS generation

Lungs were inflated using a balloon catheter with prewarmed (38–42 °C) agarose (Molecular Biology Grade, bioWORLD, Dublin, OH, USA, cat. no. 40100104-3) solution (0.8% in HBSS, Bioworld, Dublin, OH, USA), and then cooled (2–8 °C) for approximately 45 min to facilitate the agarose solution’s transition to gel. From the solid gel inflated lungs, 1–2 cm thick sections were cut using a knife and/or scalpel and placed in cold slicing buffer ^20^. The sections were cut further using a MD2300 or MD5000 low-speed coring press (Alabama Research and Development, Munford, AL, USA) to generate cylindrical cores (∼8 mm diameter). Finally, the cores were thinly sliced (400–500 µm thickness) using the Krumdieck MD4000 tissue slicer (Alabama Research and Development) according to the manufacturers’ protocols. The PCLS batches were collected and stored (<24 h) in cold slicing buffer until they underwent cryopreservation.

### PCLS cryopreservation

Using a sterile cotton swab or fine-tip forceps, up to 6 PCLS samples were carefully placed into a cryovial containing approximately 1.5 mL cryopreservation buffer (proprietary; Advanced In Vitro, LLC Frederick, MD, made at the Institute for In Vitro Sciences (IIVS), Gaithersburg, MD) and then frozen at ≤−60 °C at an approximate rate of −0.4 to −1.5 °C/minute for at least 4 and up to 72 h. The cryovials were then transferred to the vapor phase of a liquid nitrogen tank for long-term storage (Patel et al., 2023). The length of time of frozen storage prior to use in our experiments was variable across donors but within a year of procurement.

### PCLS thawing and preparation for spatial transcriptomics

Frozen samples were transported in dry ice to the Genetics and Genomics Laboratory, Children’s Hospital, Boston. They were rapidly thawed, rinsed twice in 12 mL prewarmed (37 °C) PCLS acclimation medium (Marimoutou et al., 2024), and re-frozen in optimal cutting temperature compound (TissueTek Sakura). Frozen tissues were then sectioned (12 µm sections) using a cryostat at −20 °C. Sections were transferred onto pre-chilled Visium Tissue Optimization Slides (10× Genomics, 3000394), and then to Visium Spatial Gene Expression Slides (10× Genomics, 2000233).

Spatial gene expression assay – Visium Tissue Permeabilization Optimization, Gene Expression Library Construction, and Sequencing: First, frozen tissue sections were stained for hematoxylin and eosin (H&E according to the Visium Spatial Gene Expression User Guide (10× Genomics, CG000239 rev. F). Next, the frozen sections were permeabilized on a capture area composed of a glass slide printed with an array of ∼5000 oligonucleotide 55µm spots. Permeabilization time was determined using the tissue optimization slide (10× Genomics, CG000239 rev. F). Each spot contains 82-nt-long single-stranded DNA oligonucleotides with the following features: a poly(dT) region to capture mRNA polyA tails, a 12-nt unique molecular identifier (UMI), a 16-nt spatial barcode specific for the spot, and a partial Illumina sequence. Once the cells in the tissue are permeabilized, mRNAs diffuse to the capture spots and hybridize with the oligonucleotides where they are subsequently converted to double-stranded cDNAs. The second-strand cDNA is then released from the glass slide, fragmented, indexed, amplified, and sequenced using an Illumina platform. Each cDNA contains a spatial barcode to enable the calculation of cDNA abundance at specific spatial locations in the original tissue slide upon sequencing. cDNA amplification, Quality Control, and gene expression library construction is conducted according to 10× Genomics user guide (CG000239 rev. F). The gene expression libraries were quantified using a Qubit fluorometer (Thermo Fisher Scientific) and checked for quality using DNA HS bioanalyzer chips (Agilent). Sequencing depth was calculated based on the percent capture area covered by the tissue, and the 10× Genomics recommended sequencing parameters were used to run on the Novaseq X10B (Illumina).

### 10x Genomics Visium data analysis

To exclude the spots within the transferred collagen I/cell layer that do not contain cells, H&E images were manually aligned using the Loupe browser (v6.5.0, 10x Genomics). Aligned images were then processed as follows. The sequencing data were demultiplexed using the unique sample index barcodes in both i7 and i5 index reads by the Single Cell Core Facility at the University of Copenhagen. The raw Visium data was preprocessed using SpaceRanger v2.0.1 (10x Genomics), including aligning reads to the human reference genome (GRCh38-2020-A), and generating spotxgene expression count matrices along with the corresponding spatial location matrix. Using the generated expression count matrices, an in-house Spatial transcriptomics pipeline was built based on the Seurat R package (R v4.2.3, Seurat v4.3.0) ^53,54^ including quality control, spot filtering, normalization, spot deconvolution, and visualization. Spots expressing greater than 50 genes and mitochondrial read content below 20% were considered viable. Spots of good quality were filtered, and the data was normalized using the SCTransform method in sctransform package (v0.3.5). The integration of data across various samples and conditions was then performed by employing the Harmony (v0.1.1) integration method ^55^.

### SPOTlight cell type interference

We used an integrated cell atlas of healthy and diseased lung scRNAseq data as the reference ^25^, focusing specifically on lung, lung parenchyma, and airway tissues. Level 3 cell types were used, with further filtering to exclude cell types containing fewer than 100 cells and any unknown cell types. We then used the SPOTLight method to infer cell types per spot in our spatial transcriptomics data by leveraging the reference scRNAseq datasets ^26^. The method uses non-negative matrix factorization (NMF) to estimate the cell type contributions and identify the dominant cell type by weight. This approach provides insights into cell type distribution, proportions, and tissue composition within the spatial context.

### Statistical Analysis and Data Visualization

All statistical analysis was performed using R. Significance between differentially expressed genes was determined based on the Wilcoxon rank test with Bonferroni multiple test correction (P<0.05). Significantly enriched pathways were calculated using P values with Bonferroni multiple test correction (P<0.05). Spatial plots, uniform manifold approximation and projection plots, and differential gene expression heatmaps/dot plots were all generated using Seurat. Bar plots of the cluster distributions and the deconvolution matrix were generated using ggplot2 (v3.4.1).13 Volcano plots based on differential gene expression were made using the EnhancedVolcano R package (v3.16). Dot plots of enriched pathways were generated via the dotplot function in clusterProfiler. Networks of enriched pathways were generated via the cnetplot function in the clusterProfiler Bioconductor package (4.8.3).14 Spatial plots of samples were generated using Seurat and Scanpy packages.

### DEG and pathway enrichment

For the differential gene expression analysis, we used Seurat’s FindMarkers function with default parameters, applying the Wilcoxon rank-sum test with a logfc.threshold of 0.2 and min.pct of 0.1. Additionally, differentially expressed genes (DEGs) were filtered to retain only those with an adjusted p-value lower than 0.05 (all DEG Table S1). KEGG pathways and gene ontology (GO) enrichment tests were performed by the clusterProfiler v4.6.2 R package ^56^. A pathway or GO term was treated as significantly enriched if an adjusted p-value (with Benjamini-Hochberg correction) was lower than 0.05. All bar plots and dot plots illustrating significant pathway or GO terms were created using the enrichplot v1.18.3 R package ^57^.

### Cell–Cell Communication Analysis

We used CellChat (v2.1.2) along with its provided database of ligands and receptor pairs for humans to identify patterns of cell-to-cell communication. To assess the likelihood of communication, we followed published methodologies ^45^. We applied these methods at both the ligand-receptor pair level and the pathway level. To ensure reliable and significant findings, communication between cell types observed in fewer than 10 cells with adjusted p-values greater than 0.1 was excluded. The plots were generated using the built-in functions within CellChat. Significant interactions were identified based on a statistical test that randomly permutes the group labels of cells and then recalculates the interaction probability and the signaling pathways.

## Results

### Spatial transcriptome profiling identifies cell- and gene-specific changes in fibrotic PCLS

Within cryopreserved and then thinly sliced human PCLS (Figure 1A), 23 different cell populations were identified using spatial transcriptomics. Specifically, following preprocessing and quality control (QC) to ensure data consistency and reliability (Figure1, Figure S1), we integrated oligonucleotide spots from three fibrotic patients and three controls (Figure 1B, Figure S1) into one data manifold, representing 21,213 spots. First, we normalized the dataset to eliminate expression differences caused by unavoidable technical reasons such as unequal amplification during PCR. Next, we performed batch effect correction using the approved package by 10x Genomics and also the preferred package by independent benchmark with various covariates ^24^, Harmony. Overall, such an approach allowed the successful integration of the spatial transcriptomics data, as seen in the Uniform Manifold Approximation and Projection (UMAP) graph (Figure 1C), to conduct downstream analysis. Utilizing previously published scRNA-seq datasets from human ^25^, we integrated spatial transcriptomics and scRNA-seq data using the SPOTLight method^26^. This method annotates cell types in each spot and assesses the spatial distribution of cells in the tissue by estimating the cell type compositions of each spot. SPOTLight then calculates a deconvolution weight of all cell types within each spot (Figure S2). By identifying the cell types with the highest deconvolution weight amongst the cell mixtures for each spot, this approach provides a refined understanding of cell type distribution across the tissue. Specifically, we identified 23 cell populations/topics characterized by marker gene expression profiles, including fibrosis-relevant cells (Figure 2A-B). Notably, we identified cell types historically underrepresented in scRNA-seq, including endothelial and rare cells.

**Figure 1.**
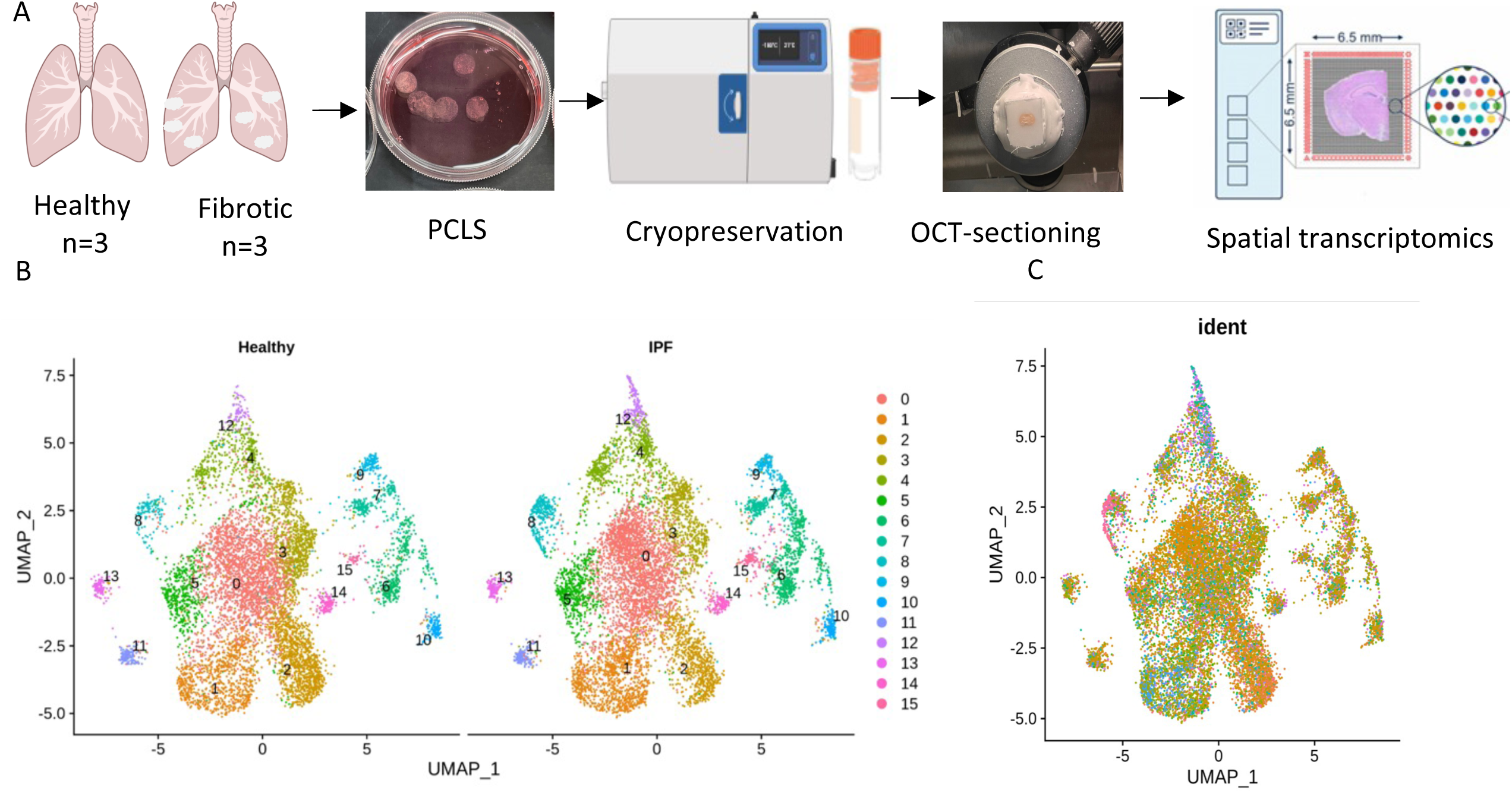
Experimental workflow and spatial transcriptomic profiling of human lung tissue from healthy and PF donors reveals clear differences in cellular composition and abundance. (A) Schematic representation of the experimental pipeline. Human PCLS were collected from healthy controls and patients with pulmonary fibrosis (PF), and cryopreserved. On the day of the experiment, they were rapidly thawed, embedded in Optical Cutting Temperature (OCT) compound, cryo-sectioned, and transferred to Visium spatial gene expression slides. The slides were processed for spatial transcriptomics using the 10x Genomics Visium platform. (B) UMAP plots show clustering of transcriptomic profiles from healthy (left) and PF (right) sections. These clusters, color-coded by cellular identity, reveal clear differences in cellular composition and abundance. (C) When combined across all tissues sections from all measurements, the integrated UMAP plot confirms integration while revealing distinct spatial transcriptomic patterns

**Figure 2.**
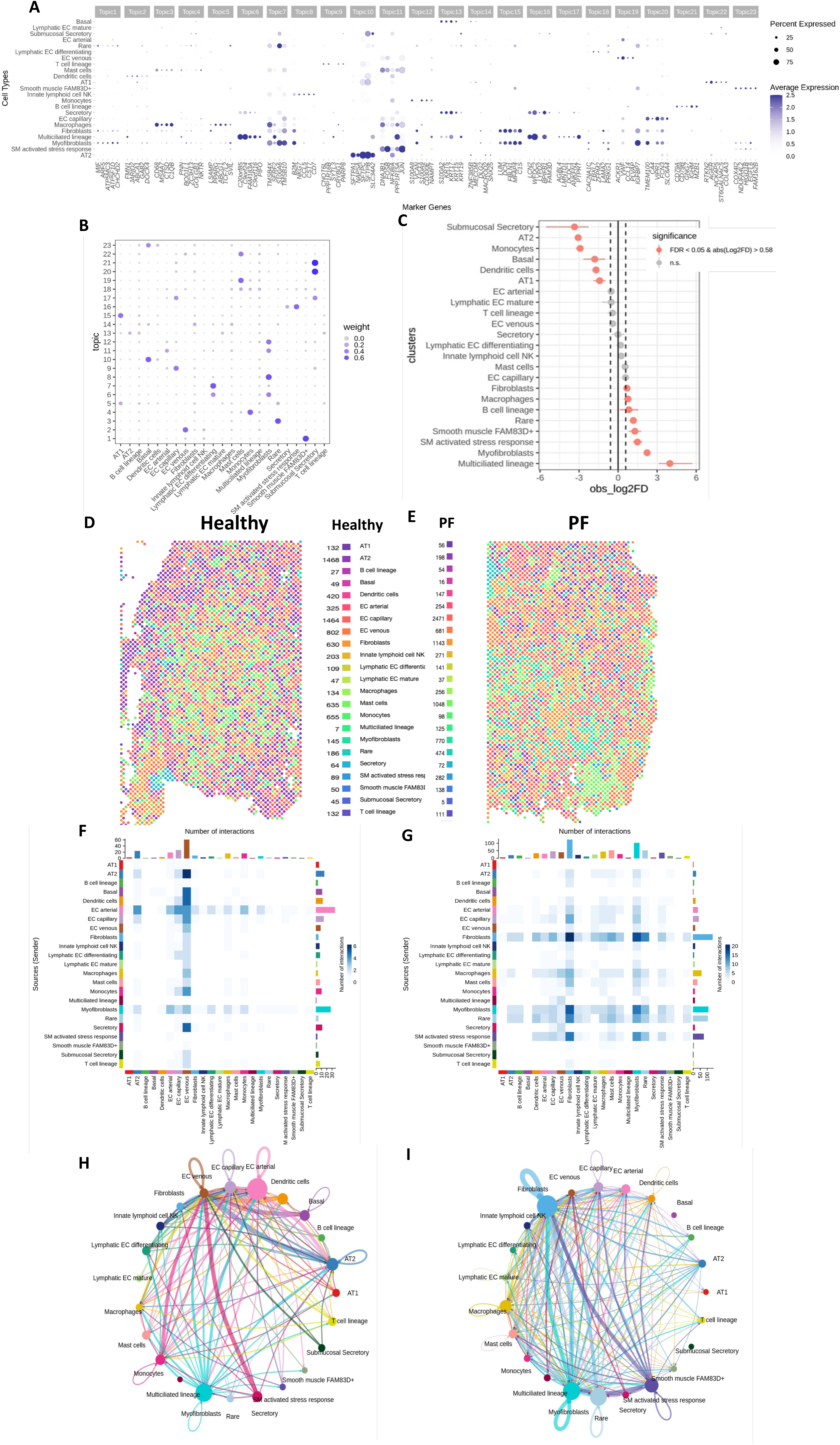
Spatial analysis reveals distinct cellular composition and unique cell-cell interactions in healthy vs PF tissue sections. (A) Gene expression across annotated cell clusters were grouped into “topics”. Dot size represents the percentage of cells expressing the gene; color intensity indicates average expression. (B) Topics were annotated based on marker gene expression. (C) scProportion test depicts differentially abundant clusters between healthy vs PF sections. Clusters enriched either in healthy or PF sections are shown in red (FDR < 0.05 and |log2FC| > 0.58). (D) Spatial maps of control lung tissue showing cell types in the region. Cell types can be visualized by cell identity (left) and annotated with cell numbers per type (right). (E) Spatial maps of PF lung tissue showing cell types in the region. Cell types can be visualized by cell identity and annotated with cell numbers per type. (F) Heatmaps of cell-cell chat analysis to represent the number of interactions between source and target cell types in healthy (left) and PF (right) sections. (H–I) Circos plots to visualize intercellular communication networks in healthy (H) and PF (I) sections. Line thickness corresponds to the number of predicted interactions; colors indicate the source cell type.

### Cell deconvolution identifies differences in cell composition

Compared to healthy PCLS, fibrotic PCLS had increased occurrence of fibroblasts, macrophages, myofibroblasts, and rare cell types including ionocytes, neuroendocrine cells, and tuft cells, in close proximity with myofibroblast (Figure 2C-E). Fibrotic PCLS also retained a surplus of smooth muscle-activated stress response-related cells and the stromal cell type, smooth muscle FAM83D^+^. In contrast, the fibrotic PCLS had decreased composition of alveolar epithelial populations such as basal, AT1 and AT2 cells, as well as dendritic cells (Figure 2C-E).

### Spatial analysis identifies differences in inter-cellular communication

Spatial interactions were markedly altered in the fibrotic PCLS, with fibroblasts, myofibroblasts, macrophages, and vascular endothelium emerging as central hubs of multi-cellular interactions (Figure 2F-I), and stromal–immune crosstalk and fibroblast signaling as dominant intercellular communication pathways. These data support the concept of a reprogrammed fibrotic niche maintained by persistent ECM remodeling, immune activation, and impaired epithelial regeneration ^27–29^. In contrast, intercellular communication in the healthy PCLS were dominated by epithelial–vascular interactions (Figure 2F, H).

### Spatial analysis identifies differences in cell-type specific gene expression patterns

By utilizing the upper and lower quartiles on the normalized data using SC Transform, we categorized the PCLS regions into ‘high’, ‘medium’, ‘low’, and ‘no expression’ of collagen and fibrotic genes (Figure 3, Figure S3A-B). Healthy PCLS displayed medium expression of collagen genes (Figure 3A, C). In contrast, fibrotic PCLS displayed high expression of collagen genes (Figure 3B, C). Similarly, while healthy PCLS displayed medium expression of fibrotic genes (Figure 3D, F), fibrotic PCLS displayed high expression of the same physical quantities (Figure 3E, F). Although the majority of the spatial regions of collagen and fibrotic gene expression were categorized as either “medium” and “low” in healthy PCLS, most of these regions were categorized as either “high” and “medium” in the fibrotic PCLS.

**Figure 3.**
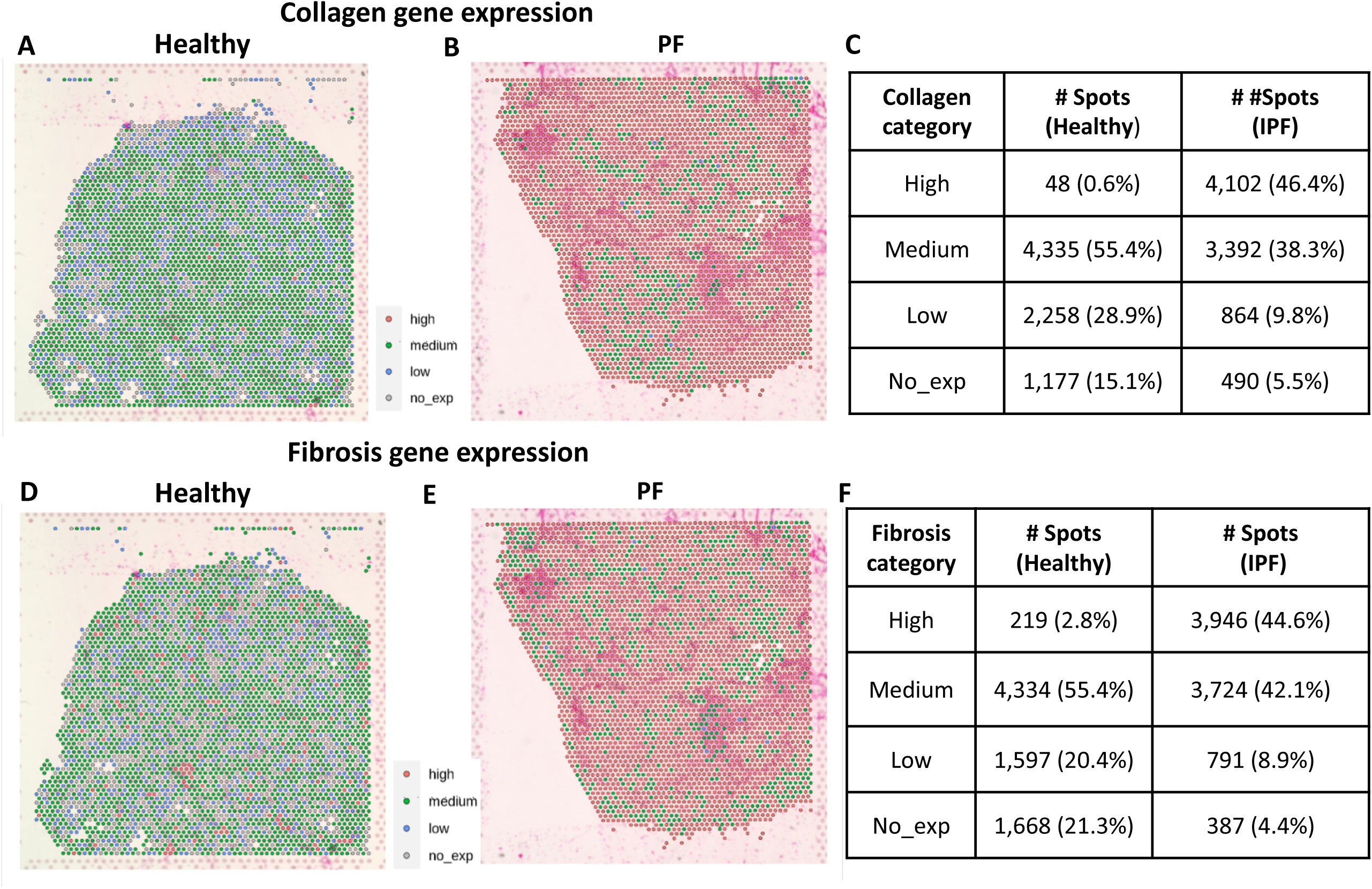
Spatial transcriptomic mapping reveals significantly greater and spatially clustered collagen and fibrotic gene expression in PF sections. (A-B) Spatial distribution of collagen gene module scores across healthy (A) and fibrotic sections (B). Spots are colored based on module score levels: high (red), medium (green), and low (blue) no expression (gray). (C & F) Corresponding quantification tables summarizing gene module scores, spot counts, and classification thresholds used in spatial analysis. (D-E) Expression of fibrosis-related genes in healthy (D) and PF sections (E).

### Spatial transcriptomics reveals fibroblast activation and niche signaling in fibrotic PCLS

Compared to healthy PCLS, fibrotic PCLS had enhanced expression of extracellular matrix (ECM)-associated genes such as *COMP, FN1, THBS2*, and multiple collagen genes (*COL6A3, COL1A2, COL3A1*) (Figure 4A). Within spatial domains of these enhanced ECM genes, we identified significant activation of three salient niche-signaling pathways: (1) the matricellular glycoprotein, *SPARC*, (2) Relaxin signaling, and (3) multiple immunoglobulin transcripts, including *IGHA1, IGHG1, IGHG3,* and *IGKC* (Figure 4B, C). While *SPARC* has previously been shown to modulate fibroblast–ECM interactions and to promote fibrotic remodeling through regulation of collagen deposition and integrin signaling pathways^30^, the relaxin signaling pathway is implicated in fibrotic alleviation via modulation of metalloproteinase activity and ECM degradation ^31^. Finally, we discovered immunoglobulin genes to be canonically expressed in plasma cells and B lymphocytes, suggesting either immune cell infiltration or co-localization with fibroblasts in immune–stromal niches. Given the 55-μm diameter capture spots of the Visium platform, this spatial co-expression likely reflects multicellular environments. This observation is also consistent with recent spatial and single-cell atlases of the human lung that identified spatially restricted fibroblast subtypes—including alveolar and adventitial fibroblasts—whose localization correlates with immune cell presence and fibrotic remodeling^25,32,33^

**Figure 4.**
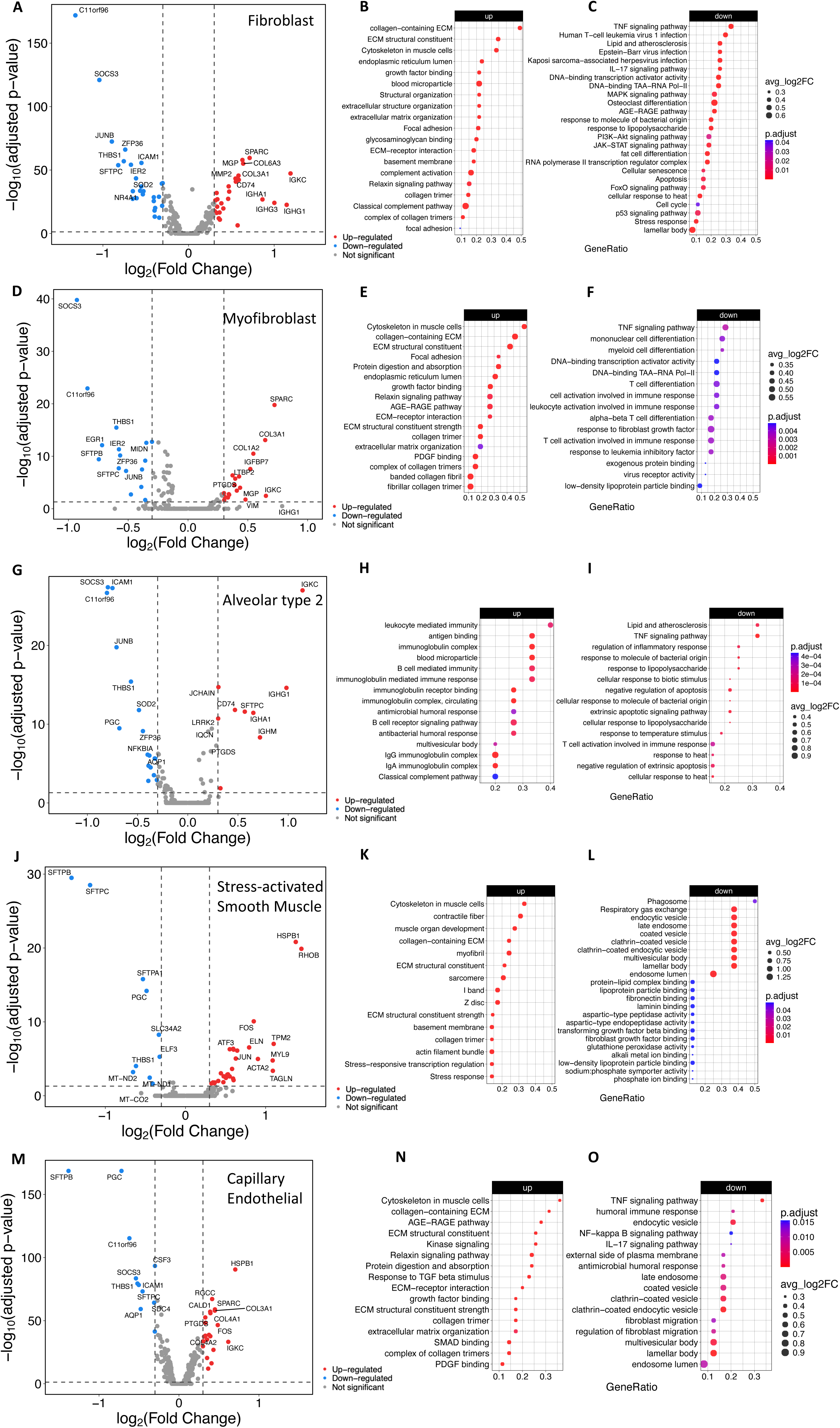
Transcriptional reprogramming and pathway enrichment reveals unique genes, pathways, and phenotypes, primarily targeted to immunomodulation, stress-activation, mechanotransduction, and ECM production in PF cells. (A–O) Differential gene expression and pathway enrichment analyses for five key stromal and endothelial cell subtypes in IPF versus healthy lung tissue: (A-C) Fibroblasts, (D-F) Myofibroblasts, (G-I) Alveolar type 2 cells, (J-L) Stress activated smooth muscle cells, and (M-O) Capillary endothelial cells. Left panels: Volcano plots showing significantly upregulated (red) and downregulated (blue) genes (FDR-adjusted p-value < 0.05, |log₂FC| > threshold). Selected genes of interest are labeled. Right panels: Dot plots depicting results of Gene Ontology (GO) and KEGG pathway enrichment analyses for upregulated (B, E, H, K, N) and downregulated (C, F, I, L, O) genes in each cell type. Dot size indicates gene ratio; color represents adjusted p-value; color intensity reflects average log₂ fold-change.

### Spatial transcriptomics reveal a stress-adapted, ECM-producing myofibroblast population in the fibrotic niche

Within fibroblast-rich regions of fibrotic lung tissue, we observed upregulation of canonical ECM constituents (*COL1A1, COL1A2, COL3A1, COL4A1, COL6A3*) and matricellular proteins (*SPARC, TIMP1, IGFBP5, LTBP2*), alongside cytoskeletal and contractile markers including VIM, ACTB, and PFN1(Figure 4D). These findings are consistent with prior spatial and single-cell studies identifying ECM-producing ACTA2⁺ myofibroblasts as central mediators of fibrosis ^32,33^. The detection of *CLDN5* and *EPAS1* in the same regions suggests concurrent endothelial remodeling or endothelial-to-mesenchymal transition (EndoMT), as has been proposed in vascular remodeling in IPF ^34^. In contrast, downregulated genes in these same spatial domains included immediate-early response regulators (*JUNB, NR4A1, GADD45B, IER3*), immune checkpoint genes (*SOCS3, ZFP36*), and epithelial markers (*SFTPB, SFTPC*), pointing to suppression of inflammatory resolution pathways and alveolar epithelial injury or absence (Figure S4). This gene pattern suggests that myofibroblasts in IPF fibrotic foci exhibit a non-resolving, stable profibrotic transcriptional state, consistent with IPF persistence ^35^. Pathway analysis of the upregulated genes revealed a highly interconnected module of ECM proteins and ECM-regulatory factors, and downregulated gene network was enriched for transcriptional repressors and immune-regulatory signals, suggesting myofibroblasts actively suppress regulatory inputs that may otherwise limit ECM production (*Figure 4E, F*).

### Within the fibrotic niche, alveolar type 2 (AT2) cells exhibit a stress-adapted, immune-modulatory state with impaired repair signaling

Not only were AT2 cells fewer in fibrotic PCLS (Figure 2C-E), the remaining cells revealed a transcriptional landscape consistent with chronic stress adaptation, immune crosstalk, and disrupted regenerative capacity. Specifically, the expression of *TSC22D3, PTGDS,* and *LRRK2,* indicates ongoing response to epithelial injury and oxidative stress^36–38^. Furthermore, induction of immunoglobulin-related genes (*IGHA1, IGHM, IGHG1, IGKC*) and immune regulatory transcripts (*CD74, JCHAIN*) suggests atypical immune–epithelial signaling within fibrotic niches (Figure 4G-I). In parallel, key injury-response and stress-regulatory genes were downregulated, including *JUNB, SOCS3, ZFP36,* and *NFKBIA*, which normally act as negative feedback regulators of inflammation. Downregulation of *SFTPB, AQP1*, and HSPA1A reflects impaired surfactant production and diminished epithelial integrity, while decreased expression of mitochondrial and metabolic genes (*PGC, SOD2, GPX3*) suggests energetic dysfunction. Multiple NF-κB inhibitors (e.g., *NFKBIA, NFKBIZ, BCL3*) were suppressed, indicating potential sensitization to chronic inflammatory stimuli. These findings support a model in which AT2 cells in fibrotic lungs remain lineage-committed but locked in a maladaptive transitional state, unable to fully regenerate alveolar epithelium or to restore homeostasis. This is consistent with prior single-cell and spatial analyses identifying a disease-associated KRT8⁺ transitional epithelial cell state that accumulates in fibrotic lung and fails to complete differentiation into functional Alveolar type 1 cells ^33,39,40^. Moreover, the elevated expression of *BMP1, SCGB3A1*, and *TSC22D3* in fibrotic AT2 cells may reflect an attempted but dysregulated reparative program, which paradoxically contributes to fibrosis through aberrant epithelial–mesenchymal and immune signaling. Together, these data position AT2 cells not as passive victims of fibrosis but as active participants in fibrotic remodeling, whose failed regenerative attempts and altered cytokine production may sustain the pro-fibrotic microenvironment.

### Smooth muscle–like stress-response cells in the fibrotic niche exhibit contractile, ECM-producing, and metabolically suppressed phenotypes

In fibrotic PCLS, smooth muscle (SM) cells activated stress responses consistent with mechanical activation, extracellular matrix (ECM) remodeling, and metabolic reprogramming (Figure 4J-L). This population upregulated a suite of contractile and myofibroblast-like genes, including *ACTA2, MYH11, TAGLN, MYL9, TPM2*, and *CALD1*, indicative of a stress-induced, activated mesenchymal state (Figure 4J). Additionally, matrix-associated genes such as *COL3A1, COL4A2, COL6A2, COL18A1*, and *SPARC* were elevated, suggesting active participation in fibrotic matrix deposition. Transcriptional regulators associated with mechanical signaling (*JUN, JUNB, FOS, EGR1, KLF2*) and anti-inflammatory adaptation (*ZFP36L1, ATF3, TSC22D3*) were also upregulated, consistent with a stress-responsive phenotype. Notably, genes such as NOTCH3, MCAM, and BTG2—implicated in vascular remodeling, mesenchymal cell plasticity, and cell cycle arrest—were also induced, suggesting this population may arise from or interact with vascular and perivascular compartments under fibrotic stress ^32,41^. Conversely, downregulated genes in this population included mitochondrial regulators (*MT-ND1, MT-ND2, PGC*), oxidative stress enzymes (*GPX3*), and surfactant-related epithelial markers (*SFTPA1, SFTPB, SFTPC, SLC34A2*), indicating a shift away from oxidative resilience and epithelial signaling. Reduced expression of *HSPA1A* and *THBS1* suggests suppressed cytoprotective and anti-inflammatory mechanisms. These changes echo prior reports of metabolically compromised fibroblast and smooth muscle–like cells in the fibrotic lung, particularly those localized to fibrotic foci^42^. Taken together, these findings define the SM activated stress response population as a contractile, ECM-producing, stress-adapted mesenchymal cell type with reduced metabolic capacity and diminished epithelial interactions. This transcriptional state is likely to contribute to tissue stiffening, mechanical feedback loops, and fibrotic progression.

### Spatial transcriptomics reveals distinct endothelial remodeling programs in the fibrotic niche

Arterial endothelial cells in fibrotic niches exhibited marked upregulation of contractile and smooth muscle–like genes, including *ACTA2, MYH11, ACTG2*, and *TAGLN*, alongside core ECM and remodeling regulators such as *COL1A1, COL3A1, SPARC*, and *CCN2* (Supplemental Figure 5A). This transcriptional program aligns with an endothelial-to-mesenchymal transition (EndoMT)-like state, previously described in vascular remodeling during fibrosis ^43^. Additional upregulated genes—*HSPB1, TSC22D3*, and *PTGDS*—suggest adaptation to mechanical stress and immunomodulatory signaling (Figure S5B, C). In contrast, downregulated transcripts, including *SFTPC, SOCS3, ICAM1*, and *ZFP36*, point to a loss of epithelial-like crosstalk and suppression of immune checkpoint pathways ^41^.

In venous endothelial cells, upregulated genes included transcriptional regulators (*JUN, JUND, FOS, FOSB*), stress-responsive molecules (*ZFP36, IGFBP7, HSPB1*), and ECM-related factors (*COL4A2, SPARC, HSPG2*) (Figure S5D). These patterns resemble those observed in perivascular inflammation and fibrotic vein remodeling in chronic lung injury ^33^. Downregulated genes again included *SFTPC, THBS1, SOCS3*, and *IL6*, as well as the matrix metalloprotease inhibitor *TIMP1*, suggesting an imbalance between matrix deposition and resolution pathways. This signature reflects a reprogrammed venous endothelium, with features of immunosuppression and loss of epithelial–endothelial communication ^44^ (Figure S5E, F).

Capillary endothelial cells, typically associated with gas exchange and alveolar maintenance, displayed a hybrid transcriptional state in IPF. Upregulated genes included ECM and mesenchymal markers (*COL1A1, COL1A2, COL6A3, BGN, LTBP2, SPARC*), transcriptional and inflammatory regulators (*FOS, FOSB, ATF3, EGR1, CCND1*), and angiocrine signaling factors such as *EGFL7* and *PTGDS*. Downregulated genes included *SFTPB, SFTPC, SFTPA1*, and *AQP1*, indicating the loss of alveolar epithelial proximity and possible capillary rarefaction. Suppression of *CXCL2, SOCS3*, and *NFKBIA* further reflects diminished immune-resolving signals, consistent with prior single-cell spatial analyses showing capillary regression and altered angiogenic tone in IPF ^25^ (Figure 4M-O).

Comparative analysis across vascular subtypes revealed conserved upregulation of *SPARC, PTGDS, TSC22D3, and IGFBP7*, indicating a shared program of matrix remodeling, prostaglandin signaling, and stress adaptation. Conversely, *SFTPC, SOCS3*, and *THBS1* were consistently downregulated, supporting the notion that vascular inflammation and repair programs are supressed across the IPF endothelium. These findings align with and expand upon recent spatial atlases of the human lung, which report zonally distinct endothelial states associated with fibrosis, immune infiltration, and tissue remodeling ^40^. Together, these spatially resolved data suggest that endothelial cells in IPF undergo subtype-specific transcriptional reprogramming involving ECM production, loss of epithelial–immune interaction, and acquisition of contractile and stress-adapted features. These changes are spatially compartmentalized and likely contribute to the regional heterogeneity of vascular remodeling in progressive fibrotic lung disease.

### Fibroblast-driven signaling dominates cellular interactions in IPF-PCLS

Through manifold learning and quantitative contrasts, CellChat identified differentially over-expressed ligands and receptors for each cell group^45^. There were 198 and 696 ligand-receptor pairs in healthy and fibrotic PCLS, respectively, categorized into 23 signaling pathways (Table 2). Notably, major interactions in the healthy PCLS are between endothelial cells (arterial and venous) and alveolar epithelial cell types II and monocytes, but in fibrotic PCLS, these connections are altered and become a fibroblast-mediated connection, including fibroblasts connecting to myofibroblasts, and fibroblasts connecting to rare cells. Remarkably, in the fibrotic PCLS, fibroblasts also expressed the largest number of ligands and receptors among all cell types and interacted with myofibroblast, rare, macrophages, and stress-activated smooth muscle cell types. Finally, the fibrotic PCLS was characterized with a high level of outgoing signals (ligands) corresponding to immune cell recruitment enriched pathways (*COMPLEMENT, CXCL, CCL, MIF, TNF, VISFATIN, GALECTIN*), cell growth and proliferation (*PDGF, VEGF, IGFBP, MIF, MK*), tissue remodeling (*GRN, CALCR, PLAU, ANGPTL, TWEAK*).

**Table 2.**
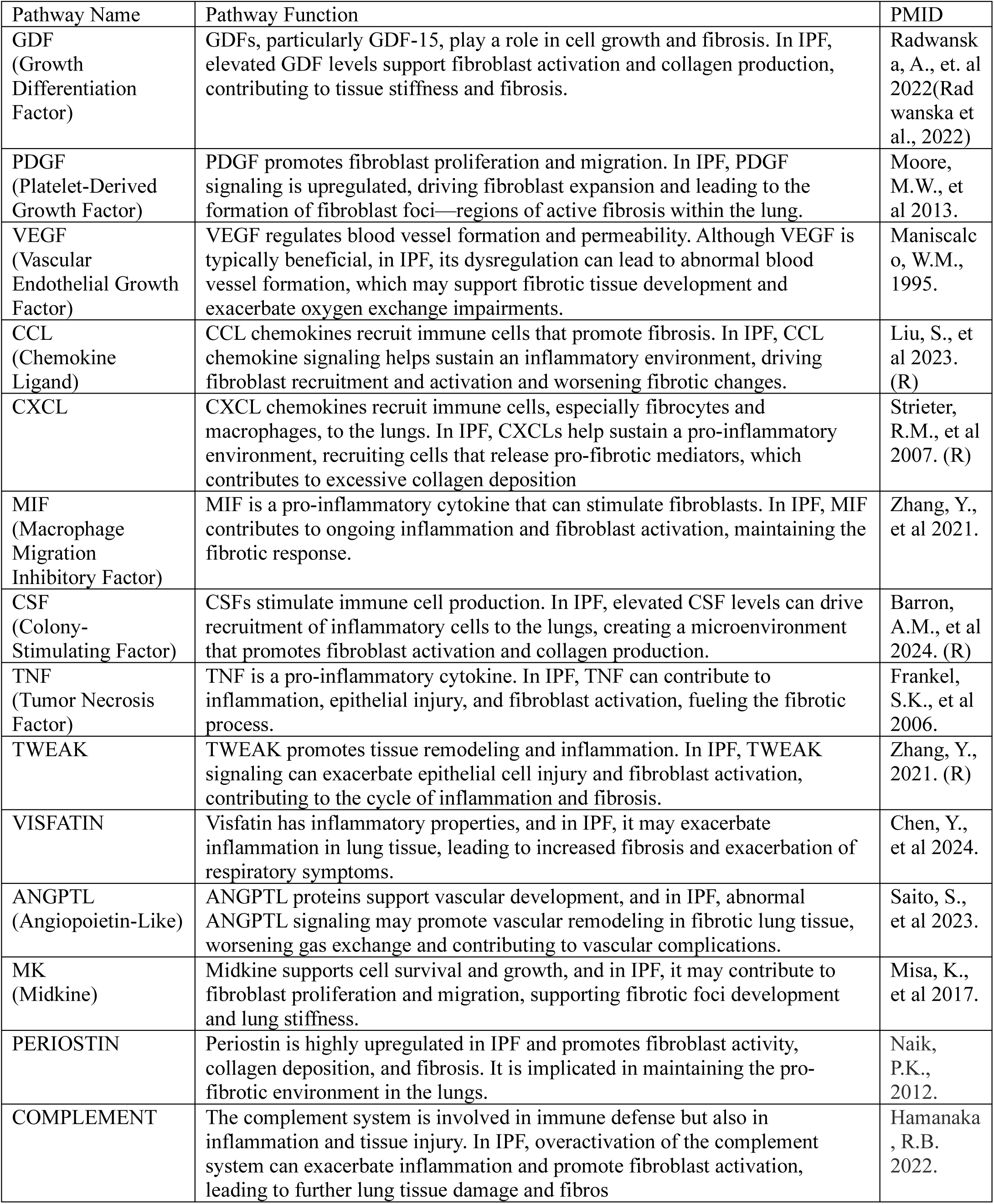
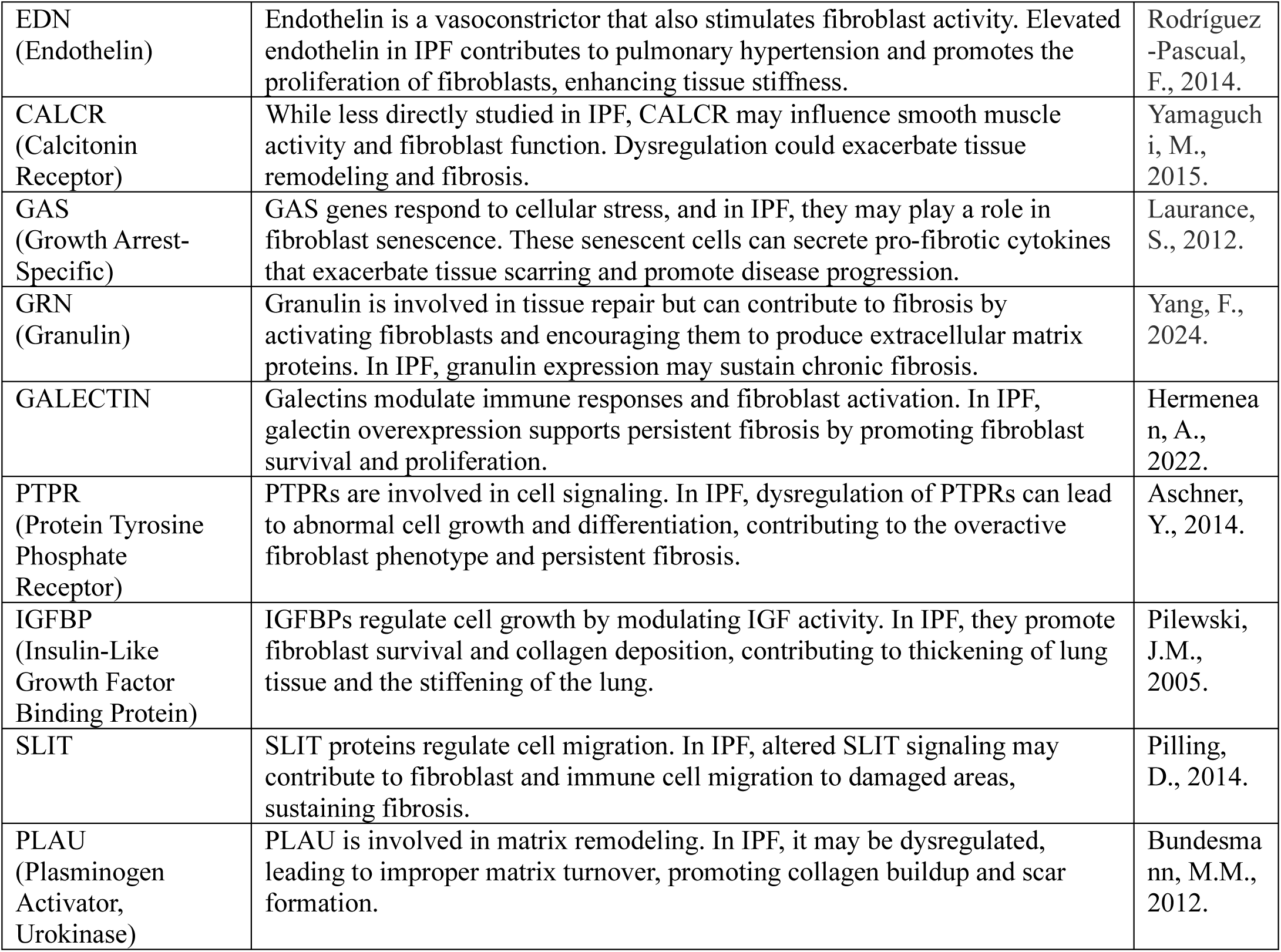
23 signaling pathways identified from ligand-receptor pairs in healthy and fibrotic PCLS.

### CXCL chemokine signaling is rewired in IPF lung with enhanced CXCL12–CXCR4 interactions and expanded fibroblast-endothelial crosstalk

In healthy PCLS, the dominant ligand-receptor pairs were largely restricted to interactions involving *CXCL1–3, CXCL6*, and *CXCL8* binding to *ACKR1*, a non-signaling scavenger receptor expressed predominantly on endothelial cells (Figure 5A). These interactions were spatially confined and primarily involved immune-endothelial regulation, consistent with homeostatic immune surveillance and vascular quiescence. In contrast, CXCL12–CXCR4 was the dominant ligand-receptor pair in the fibrotic PCLS (Figure 5B), with markedly elevated relative contributions. This shift from ACKR1-dominant interactions in the healthy PCLS to CXCL12– CXCR4 dominant interactions in the fibrotic PCLS signifies a functional repurposing of CXCL signaling toward pro-fibrotic recruitment and stromal activation. CXCL12–CXCR4 signaling has been implicated in fibrocyte and monocyte recruitment and is known to promote fibroblast migration and survival in fibrotic contexts^46,47^

**Figure 5.**
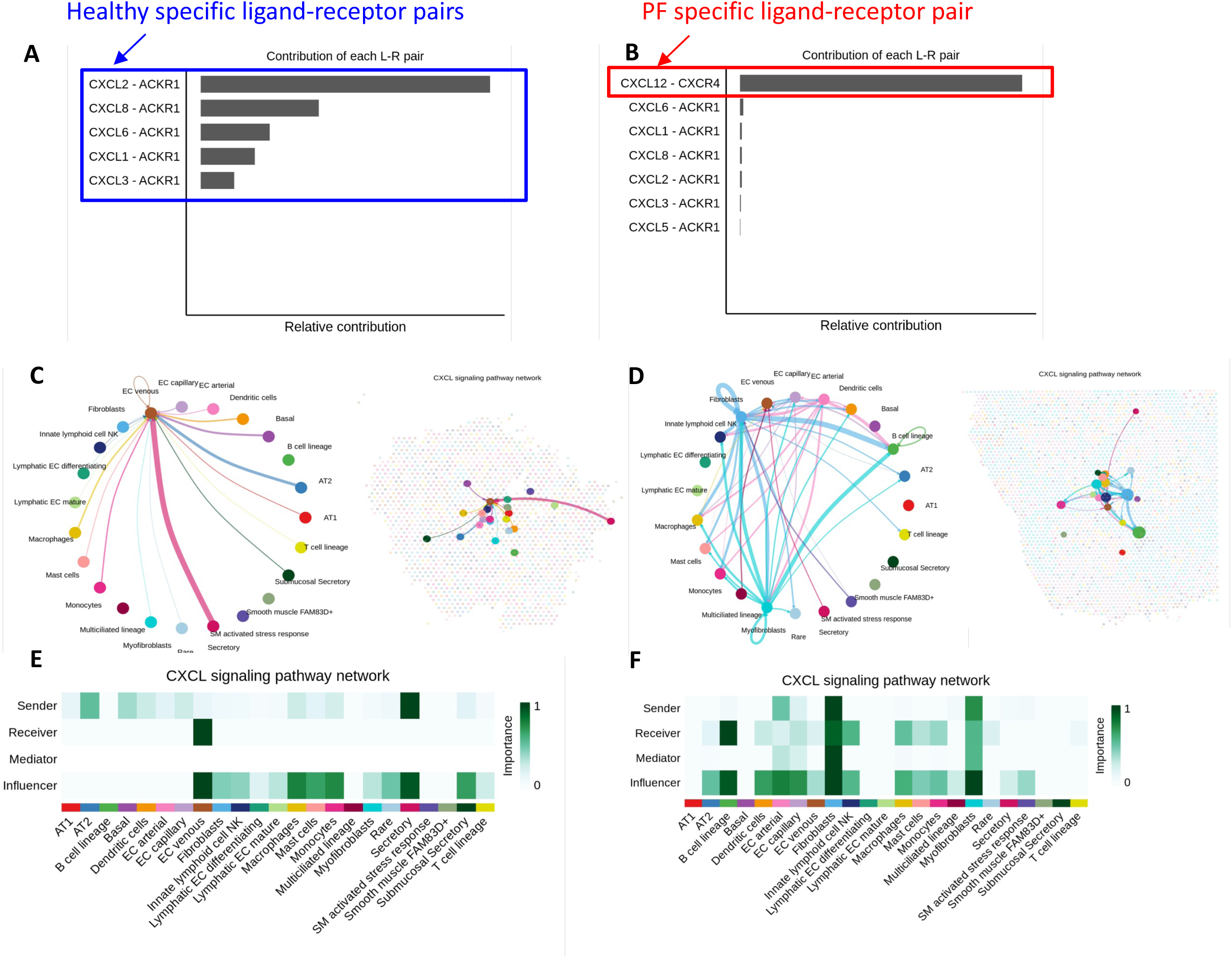
CXCL chemokine signaling is reprogrammed in PF tissues with enhanced CXCL12-CXCR4 interactions and expanded fibroblast-endothelial crosstalk. (A) Bar plot showing the contribution of individual ligand–receptor (L–R) pairs to CXCL signaling in healthy lungs. CXCL2–ACKR1 and CXCL8– ACKR1 represent dominant L–R interactions, unique to healthy tissues. (B) Bar plot highlighting IPF-specific CXCL interactions. The CXCL12–CXCR4 pair demonstrates the strongest contribution to CXCL signaling in fibrotic lungs, suggesting a shift in chemokine communication dynamics. (C–D) Circos plots visualizing cell–cell interactions mediated by CXCL signaling in healthy (C) and PF (D) sections. Arrows represent directionality from sender to receiver cell types, with edge thickness proportional to interaction strength. (E–F) Heatmaps of CXCL signaling pathway networks showing quantitative sender-to-receiver interactions between cell types in healthy (E) and PF (F) lungs. Row and column labels correspond to annotated cell clusters; color intensity reflects interaction magnitude.

Network diagrams of cell–cell communication in the fibrotic PCLS revealed that the CXCL12– CXCR4 signaling originated predominantly from capillary and venous endothelial cells, targeting fibroblasts, macrophages, myofibroblasts, and cells within the SM activated stress response cluster. Notably, these same recipient populations were less prominent or absent in the healthy PCLS, underscoring disease-specific recruitment and amplification of fibrotic crosstalk (Figure 5C-D). The CXCL signaling pathway was especially enriched in immune-modulatory fibroblast and mesenchymal populations, highlighting their dual roles as signal mediators and influencers in the fibrotic niche. These changes can contribute to excessive extracellular matrix (ECM) deposition, immune cell honing, especially macrophages and B-cells, chronic inflammation, and the further release of profibrotic cytokines, perpetuating tissue damage and fibrosis ^48^. UMAP projections confirmed the spatial localization of these CXCL networks to fibrotic regions, with CXCL12–CXCR4 signaling nodes coalescing around fibroblast-rich and immune cell–dense domains. Heatmaps of importance scores for sender, receiver, mediator, and influencer roles further emphasized the expanded intercellular influence of fibroblasts, myofibroblasts, and endothelial cells in the IPF lung, in contrast to a more restricted immune– endothelial interface in healthy tissue (Figure 5E-F). Together, these results demonstrate that CXCL signaling is rewired in fibrotic lungs, shifting from homeostatic immune-endothelial interactions to pro-fibrotic mesenchymal–immune–vascular crosstalk centered on the CXCL12– CXCR4 axis. This ligand–receptor pathway may serve as a therapeutic target to disrupt pathological cell recruitment and stromal activation in fibrotic lung disease.

### ANGPTL signaling is reprogrammed in fibrotic PCLS, shifting from homeostatic endothelial interactions to integrin-driven mesenchymal targeting

In healthy lung, ANGPTL4-mediated signaling was limited and focused on vascular-endothelial homeostasis, with top ligand–receptor pairs including ANGPTL4–SDC4 and ANGPTL4–CDH5, reflecting interaction with endothelial glycocalyx and junctional complexes (Figure 6A). In contrast, fibrotic PCLS exhibited expanded and rewired ANGPTL signaling networks (Figure 6B). ANGPTL2 interacts with the integrin receptor complex formed by ITGA5 and ITGB1 (α5β1 integrin), which plays a key role in cellular adhesion, migration, and inflammation ^49^. This interaction supports cell adhesion to the extracellular matrix (ECM) and facilitates cell migration along with release of proinflammatory cytokines and chemokines^50^. ANGPTL2 can also influence angiogenesis through α5β1 integrin, promoting new blood vessel formation in fibrotic tissue^51^. While this may be a part of normal wound healing, in the fibrotic lung, the new vasculature can support fibrotic tissue expansion and sustain fibroblast and immune cell activity. The most prominent fibrotic-specific interaction was ANGPTL2–(ITGA5+ITGB1), a receptor complex known to facilitate fibroblast adhesion, migration, and activation under mechanical and inflammatory stress. Additional ANGPTL4-based interactions with integrin components (CDH11, ITGA5+ITGB1) and matrix-interacting proteins (SDC3, SDC2) were also upregulated, suggesting a shift from endothelial maintenance to mesenchymal targeting and ECM engagement. Further analysis of cell-cell network maps in the fibrotic PCLS revealed that ANGPTL ligands were primarily secreted by fibroblasts, EC capillary/arterial cells, and SM-like stress-responsive cells, targeting a broad range of recipient populations including myofibroblasts, macrophages, and lymphatic ECs (Fig. 6C-D). This contrasted sharply with healthy networks, where signaling was spatially and functionally restricted to endothelial–epithelial crosstalk. UMAP projections of pathway activation confirmed the localization of ANGPTL-driven communication to fibrotic niches, where mesenchymal cells predominated. Quantitative heatmaps of signaling roles further illustrated that myofibroblasts and fibroblasts functioned as dominant receivers and influencers of ANGPTL signaling in fibrotic PCLS (Figure 6 E-F), whereas these roles were negligible in healthy PCLS. These findings point to a disease-associated reprogramming of ANGPTL signaling, in which fibroblasts and vascular subtypes become both sources and targets of integrin- and syndecan-mediated profibrotic cues. Collectively, these results identify ANGPTL–integrin interactions as a critical axis of fibroblast activation and intercellular coordination in fibrotic lungs, with potential roles in matrix remodeling, mechanical sensing, and immune modulation.

**Figure 6.**
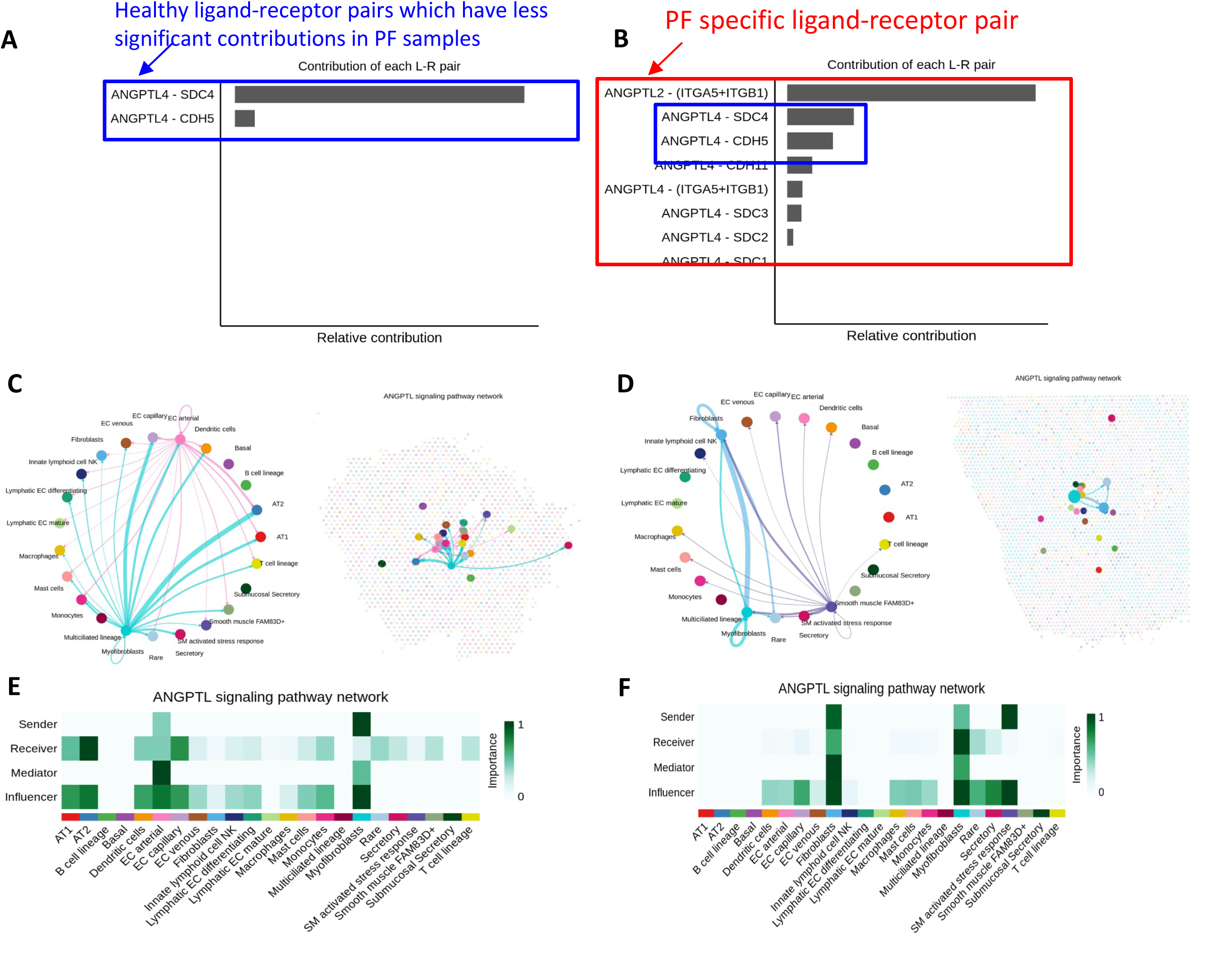
Differential ANGPTL4 ligand-receptor signaling is reprogrammed in PF tissues, shifting from endothelial interactions to integrin-driven mesenchymal targets. (A) Bar plot showing the relative contribution of individual ligand-receptor pairs within the ANGPTL4 signaling pathway in healthy tissues. ANGPTL4–SDC4 and ANGPTL4–CDH5 represent top ligand-receptor interactions in healthy tissue with minimal contribution in PF tissues. (B) Bar plot displaying the most prominent ANGPTL4-related ligand-receptor pairs in PF lungs. ANGPTL4–ITGA5+ITGB1 and ANGPTL4–SDC2 interactions are significantly enriched in PF tissue. (C–D) Network plots showing inferred ANGPTL4 signaling interactions among cell types in healthy (C) and PF (D) sections. Arrows indicate directionality from sender (ligand-expressing) to receiver (receptor-expressing) cells. Node size reflects overall signaling strength. (E–F) Heatmaps summarizing the inferred influence of sender cell types (rows) on receiver cell types (columns) through the ANGPTL4 pathway in healthy (E) and PF (F) lung sections. Color intensity indicates interaction strength.

## Discussion

Using spatial transcriptomic analysis, we have verified that the cryopreserved-thawed human fibrotic PCLS preserves cellular diversity, fibroblast and myofibroblast activation, epithelial depletion and reprogramming, and endothelial cell remodeling commensurate to never-frozen fibrotic tissue biopsies. Furthermore, fibrotic gene expression, cellular signaling networks, intercellular communication, especially fibroblast-immune cell crosstalk, and ECM deposition were co-localized to spatially defined regions called fibrotic niches.

Specifically, fibrotic niches were defined by ECM-producing, stress-adapted myofibroblasts in collagen-rich regions, characterized by elevated expression of canonical matrix genes, matricellular proteins, and cytoskeletal markers. These transcriptional signatures align with prior single-cell and spatial atlases from never-frozen PF lung explants, confirming that the cryopreserved-thawed PCLS, (using the specific methodology employed for these studies) patho-physiologically retain fibrotic mesenchymal activity. Notably, our analysis uncovered a novel result, strong co-enrichment of Relaxin signaling components in these fibrotic niches, suggesting an endogenous counter-regulatory antifibrotic response. Relaxin, via RXFP1-mediated pathways, is known to inhibit TGF-β signaling, promote MMP activation, and suppress fibroblast contractility ^31,52^. Our findings suggest that endogenous Relaxin signaling is active but unable to overcome dominant profibrotic cues, raising the possibility that exogenous Relaxin agonists may selectively benefit patients with defined spatial patterns of matrix accumulation.

Within fibrotic niches, we have also observed marked depletion of AT2 cells and associated surfactant genes (*SFTPB, SFTPC, SFTPA1*). We emphasize that this epithelial cell and gene loss is not uniform but spatially confined to the fibrotic niches. When taken together with co-localization of fibroblast expansion and immune infiltration, our findings support the emerging concept of a disease-associated transitional epithelial state that fails to regenerate functional alveoli ^33,39^. Furthermore, downregulation of immune regulatory genes (*SOCS3, ZFP36*) and mitochondrial stress responses (*SOD2, GPX3*) in these niches further reflects an energetically compromised, pro-inflammatory epithelial compartment. These findings extend prior work by showing that epithelial injury and fibroblast activation co-localize in specific spatial zones, reinforcing a model in which fibrosis is sustained by local failure of epithelial repair and dysregulated immune-stromal crosstalk.

Our data also illuminate vascular heterogeneity and reprogramming in IPF, with arterial and venous endothelial cells acquiring contractile and matrix-remodeling phenotypes, including upregulation of *ACTA2, MYH11*, and *SPARC*. The detection of EndoMT-associated markers, including EPAS1, in vascular-rich fibrotic areas, supports recent evidence that the endothelium contributes actively to fibrotic remodeling^43,44^

Furthermore, CXCL12–CXCR4 signaling, classically involved in stem cell recruitment and immune migration, was prominently rewired in the fibrotic PCLS, promoting expanded interactions between endothelial cells and mesenchymal populations. This shift from homeostatic endothelial–immune interactions to profibrotic vascular–stromal crosstalk underscores the evolving role of vasculature in PF pathogenesis and provides a rationale for targeting the vascular-chemokine axes in resolving PF.

Taken together, the preservation of these transcriptional and spatial phenotypes in the cryopreserved PCLS—with qualitative similarity to never-frozen human tissue—demonstrates the potential of such cryopreserved PCLS for interventional studies, including spatially resolved drug testing, genetic manipulation, and real-time evaluation of ECM dynamics. The platform supports high-throughput and temporally controlled experimentation, enabling multi-condition comparisons across spatial compartments that are difficult to access in vivo or in fresh human specimens. This model also opens a window into spatial biomarker discovery, identifying niche-specific gene modules (e.g., *SPARC, PTGDS, ANGPTL2*) that may serve as readouts of disease progression or therapeutic efficacy.

We acknowledge several study limitations. First, end-stage fibrotic lungs are, by themselves, insufficient to discern early stage cellular and molecular pathways of fibrotic onset and progression. Second, our acute measurements (i.e. immediately after PCLS thaw) may not be fully representative of tissue preservation during chronic fibrotic modeling (i.e. extended PCLS cultures over multiple days). Third, we are limited to the current spatial resolution of the Visium technology. Fourth, while spatial transcriptomic analysis is powerful and highly insightful, it is limited by expense and low throughput, and thus the small sample numbers considered here are a limiting factor in determining DEGs and in determining cell composition. Finally, the spatial resolution of the Visium technology is limited to spot detection and, thus, cannot truly provide single-cell resolution. The latest Xenium can now provide single-cell resolution. Future studies using newer technologies, additional time points of disease onset and development, especially in the additional presence of candidate drugs, will provide novel insights into single-cell mechanisms, fibrotic niches, drug activity, and importantly, their inter-relationships.

In conclusion, spatial transcriptomic measurements validate the applied cryopreserved-thawed cryopreservation test system as a biologically accurate and spatially informative model. Beyond basic discovery, our approach is widely applicable to drug discovery, drug validation, and insights into spatially defined and cell-type specific molecular mechanisms of drug function. Future studies employing higher-resolution platforms (e.g., Xenium, MERFISH) and perturbational assays will further expand the utility of this approach for translational fibrosis research and precision therapeutics.

## Author Contributions

R.K., I.S., and S.C., conceived and designed research; R.K., I.S., E.M, D.S, and S.C. performed experiments, analyzed data, and interpreted results of experiments; Z.A analyzed data and generated figures; R.K., and S.C. prepared figures; R.K. and S.C. drafted manuscript, R.K., M.A., J.K, Z.A., E.M., D.S., C.C., H.B., M.P.L., B.S. and S.C. edited and revised manuscript.

## Acknowledgments

This work was supported, in part, by an Alternatives Research and Development Foundation Grant (R.K.), and an NIH SBIR grant 1R43HL176301-01 (R.K., J.K, and B.S). S.C. is supported by NHLBI (R01HL152063), NINDS (1UG3NS132144), and the Cell Discovery Network, a collaborative initiative funded by The Manton Foundation and The Warren Alpert Foundation at Boston Children’s Hospital. The analysis was performed with the computational resources provided by the Research Computing Group at Boston Children’s Hospital and Harvard Medical School (Boston, MA), including High-Performance Computing Clusters Enkefalos 2 (E2), and the BioGrids scientific software made available for data analysis. We thank Dr. Rachel Knipe, Assistant Professor at Massachusetts General Hospital (Boston, USA) for helpful discussions.

---

**Supplemental Table 1: Differential gene expression in cell-specific manner between healthy and PF human PCLS.**

**Supplemental Figure S1. Spatial transcriptomics quality control metrics across multiple tissue sections.** Each row represents a different tissue section analyzed using spatial transcriptomics. For each sample, spatial feature plots (left and right images) show the spatial distribution of total UMI counts (nCount_Spatial) per spot across the tissue. The central panels display violin plots summarizing QC metrics: total UMI counts (nCount_Spatial), number of detected genes (nFeature_Spatial), and percentage of mitochondrial gene expression (percent_mito) across spatial spots. These metrics are used to assess data quality and guide filtering for downstream analysis. Color gradients on the spatial maps reflect the magnitude of nCount_Spatial, with warmer colors indicating higher transcript counts. Spots with less than 50 features, and high content of mitochondrial reads, i.e. <20% across healthy and IPF PCLS.

**Supplemental Figure S2. Spatial mapping of annotated cell types in human lung tissue sections.** SPOTLight cell type annotation proportion pie chart showing annotated cell clusters across the healthy and PF PCLS.Spatial transcriptomics maps for six lung samples (Lung_25, Lung_62, Lung_23, Lung_82, Lung_7, and Lung_57) display the spatial distribution of transcriptionally defined cell types. Each colored spot represents a spatial barcode assigned to one of 25 cell types based on transcriptomic signatures. Cell types include epithelial, endothelial, immune, and stromal populations, as well as rare and lineage-specific subsets. Notable populations include alveolar type 2 (AT2) cells, multiple endothelial subtypes (e.g., EC capillary, EC arterial), fibroblasts, macrophages, and multiciliated lineage cells. The spatial heterogeneity and organization of cell types reflect the anatomical and functional complexity of the lung microenvironment.

**Supplemental Figure 3 A. Spatial transcriptomic mapping of collagen gene expression in PCLS.** Spatial distribution of collagen gene module scores across healthy and fibrotic PCLS. Spots are colored based on module score levels: high (red) low (blue) no expression (gray).

**Supplemental Figure 3B. Spatial transcriptomic mapping of fibrotic gene expression in PCLS.** Spatial distribution of collagen gene module scores across healthy and fibrotic PCLS. Spots are colored based on module score levels: high (red) low (blue) no expression (gray).

**Supplemental Figure 4. Representative PCLS with H&E and Immunofluorescence staining.** (A–B) Representative H&E staining of control and PF PCLS. (C-D) Representative PCLS with cell type annotation on the tissue sections. (E-J) Immunostaining of surfactant proteins (SFTPC, SFTPB), NF-κB, and smooth muscle actin (SMA), along with nuclear counterstaining (DAPI), in healthy and pulmonary fibrosis (PF) in PCLS. Loss of SFTPC/SFTPB and decrease in NF-κB expression are evident in fibrotic regions, accompanied by increased SMA expression, indicating myofibroblast accumulation. (E-F) SFTPC (green), SMA (red), and DAPI (blue) in healthy (E) and PF (F) tissue. (G-H) SFTPB (green), SMA (red), and DAPI (blue) in healthy (G) and PF (H) tissue. (I–J) NF-κB (green), SMA (red), and DAPI (blue) in healthy (I) and PF (J) tissue.

**Supplemental Figure 5. Transcriptional reprogramming and pathway enrichment reveals unique genes, pathways, and phenotypes, primarily targeted to immunomodulation, stress-activation, mechanotransduction, and ECM production in PF cells. (A–F)** Differential gene expression and pathway enrichment analyses for five key stromal and endothelial cell subtypes in IPF versus healthy lung tissue: (A) Arterial Endothelial Cells, (B) Venous Endothelial Cells

