## Supplementary figures and images for "Resolving the molecular niche of pulmonary fibrosis using cryopreserved human precision cut lung slices"

### Fig.S1

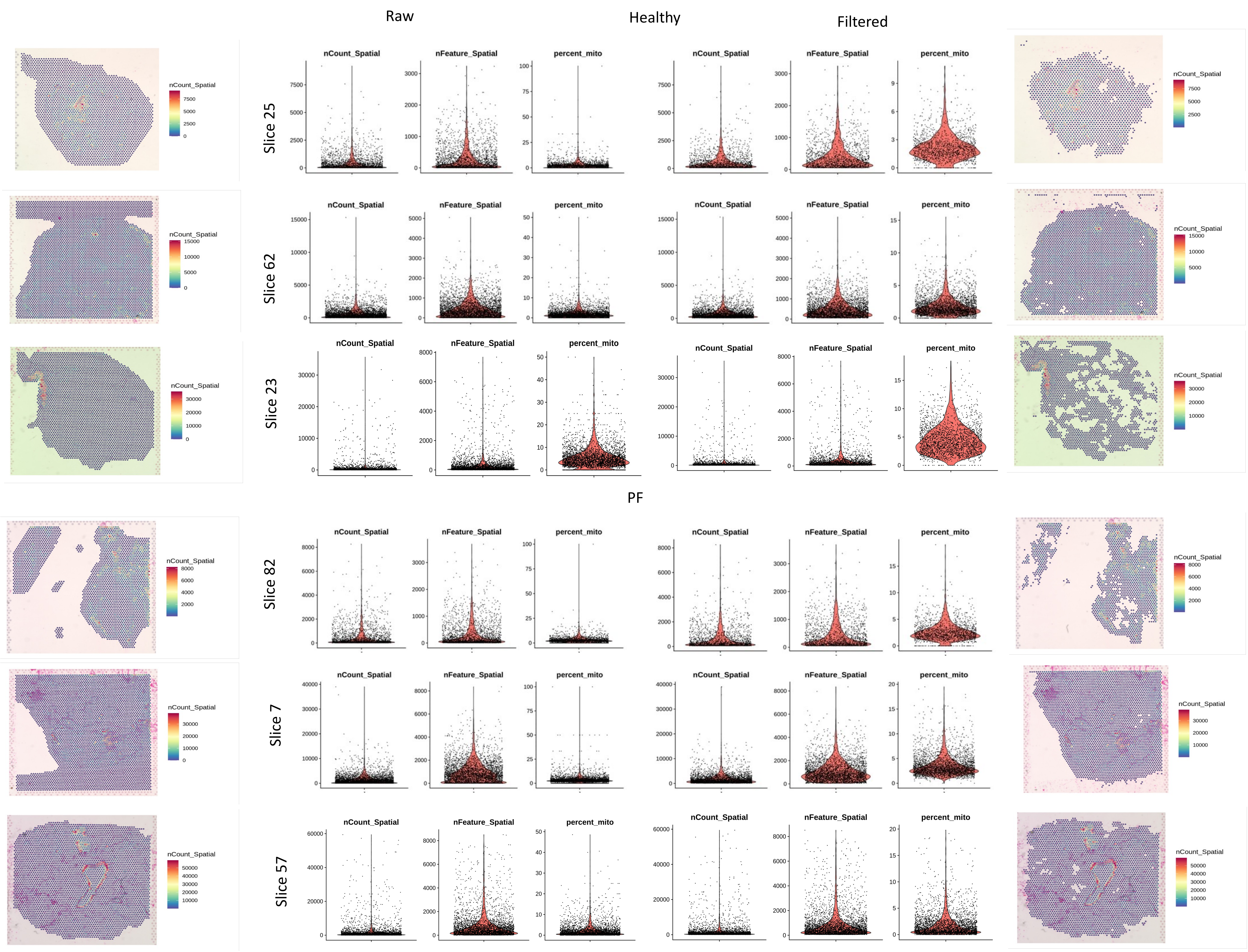

### Fig.S2

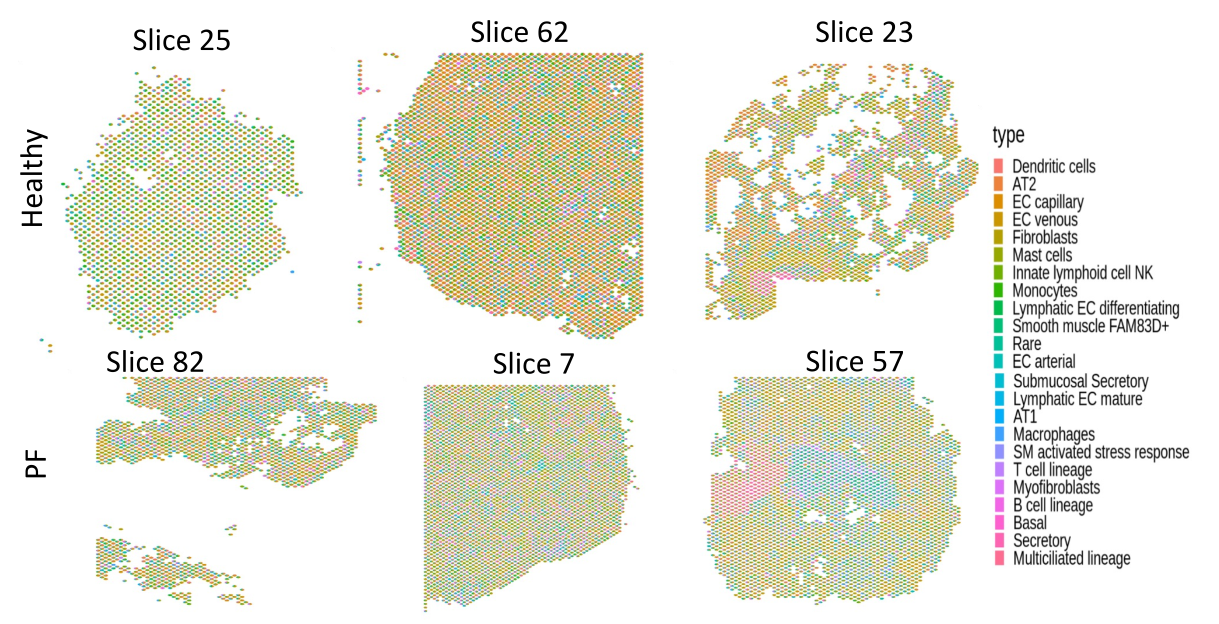

### Fig.S3A

#### Collagen genes expresion

Healthy PF

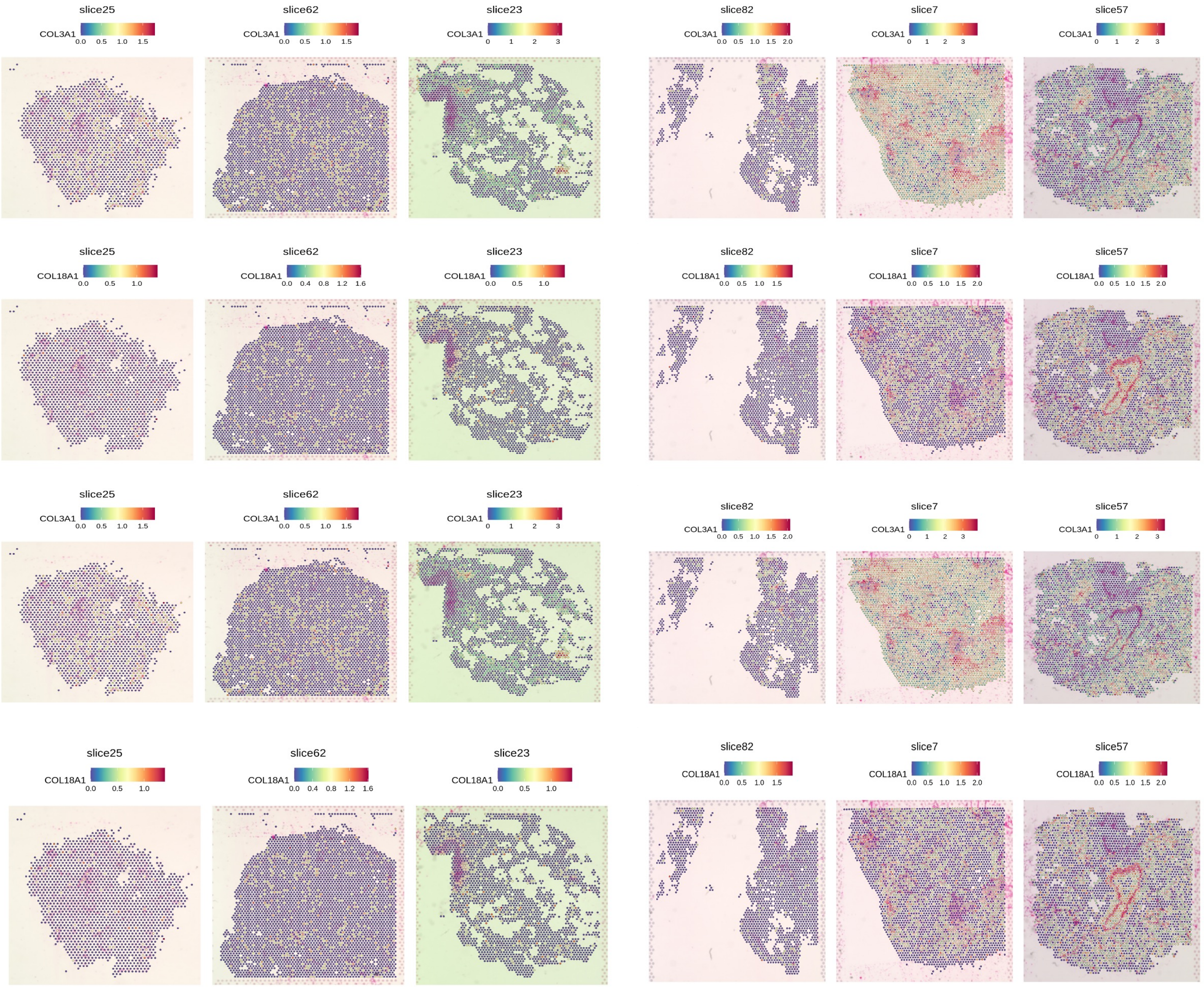


##### Total collagen gene expression

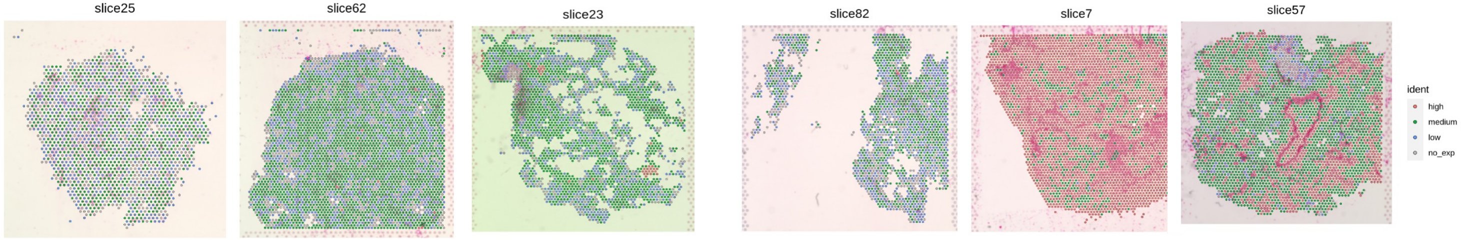

### Fig.S3B

#### Fibrosis gene expression

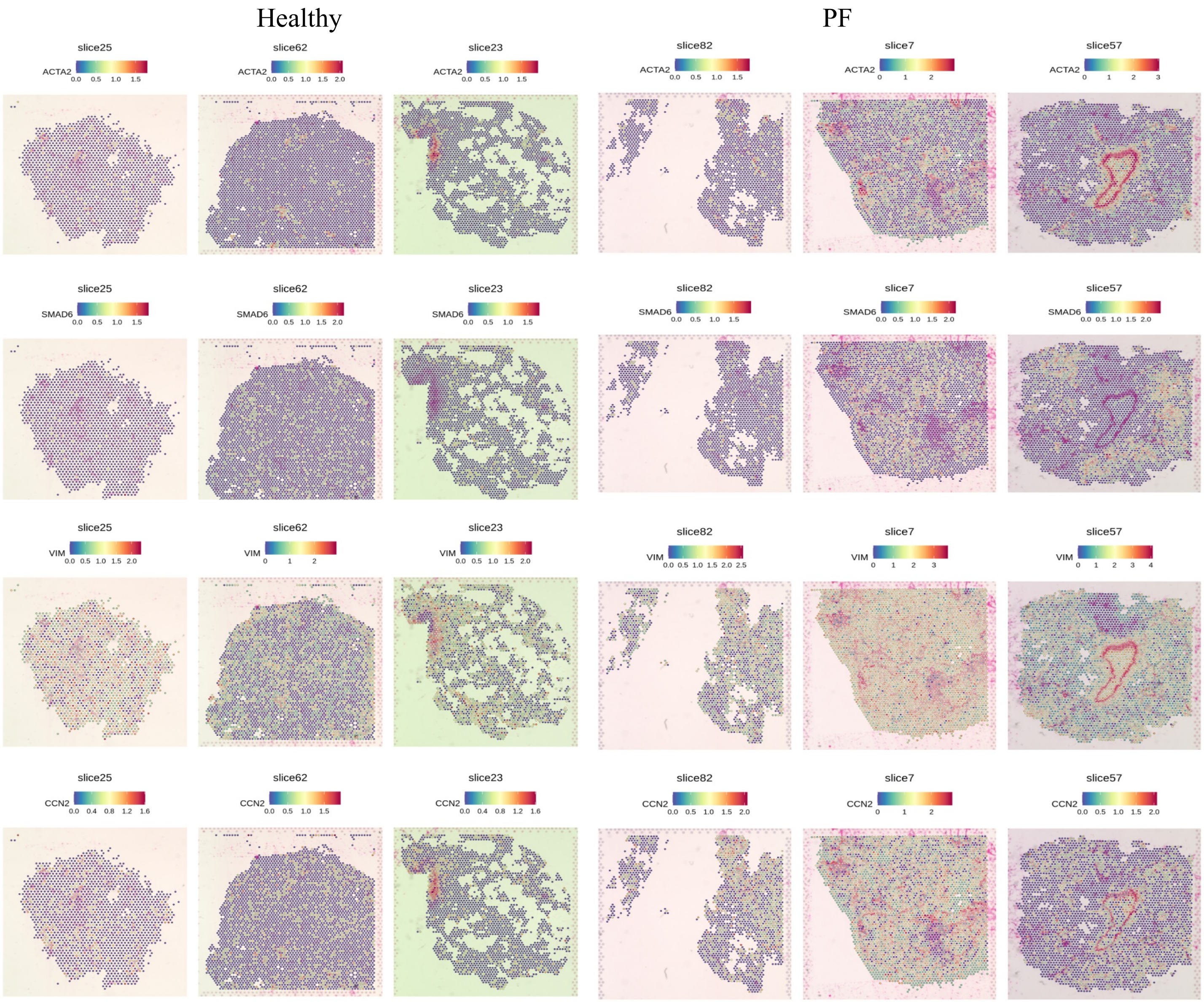


##### Total Fibrotic gene expression

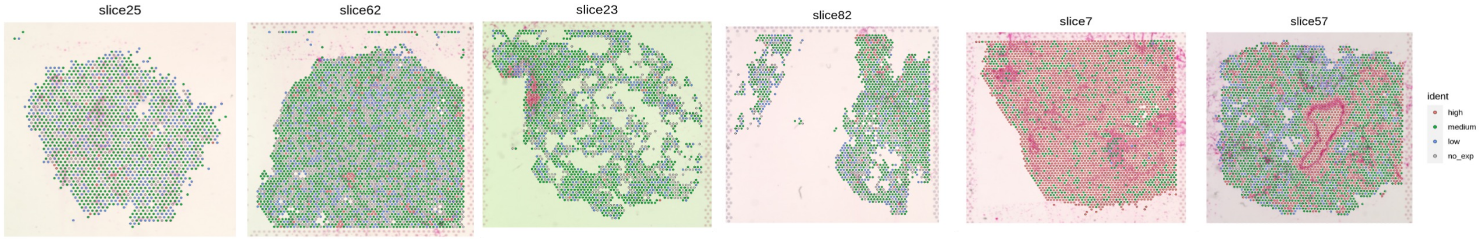

### Fig.S4

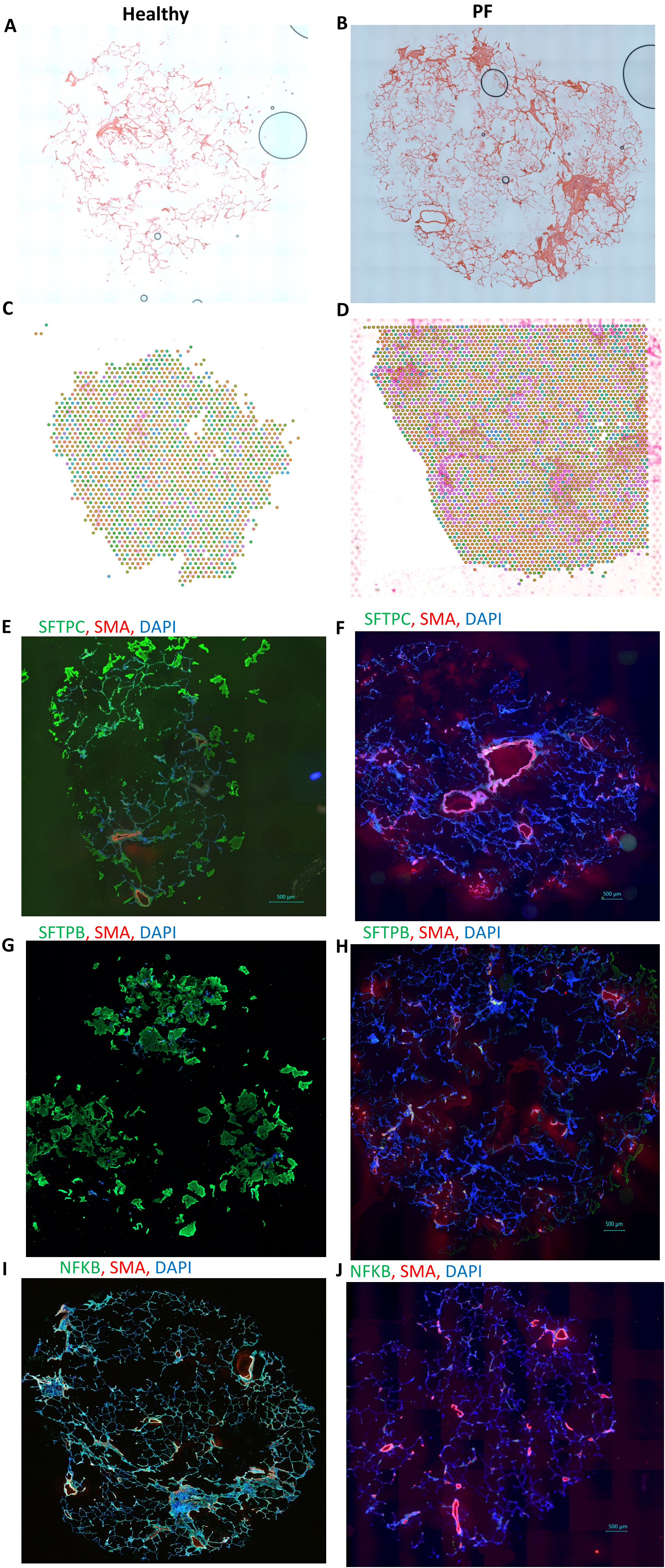

### Fig.S5

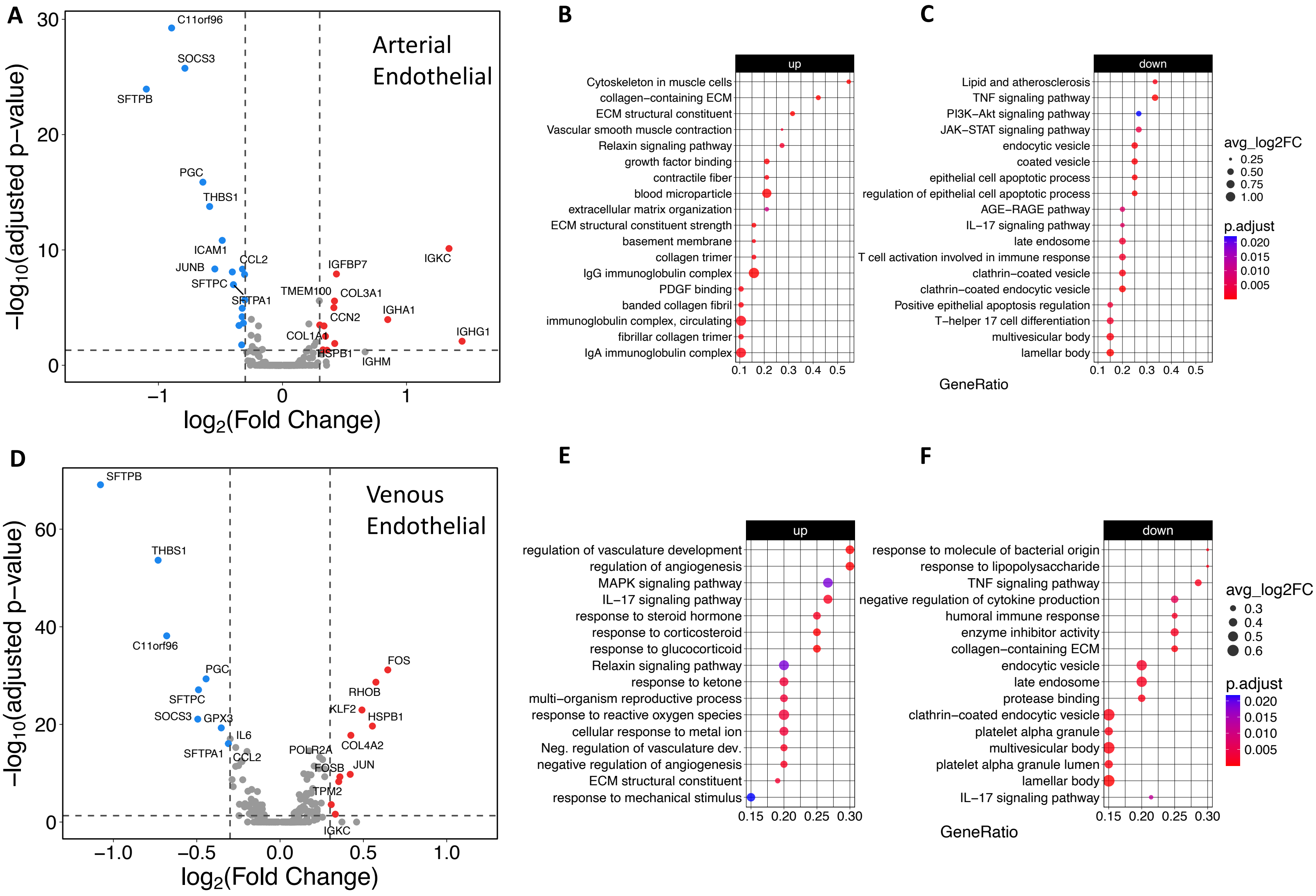
